# Tunable self-cleaving ribozymes for modulating gene expression in eukaryotic systems

**DOI:** 10.1101/739326

**Authors:** Thomas Jacobsen, Gloria Yi, Hadel Al Asafen, Ashley A. Jermusyk, Chase L. Beisel, Gregory T. Reeves

## Abstract

Advancements in the field of synthetic biology have been possible due to the development of genetic tools that are able to regulate gene expression. However, the current toolbox of gene regulatory tools for eukaryotic systems have been outpaced by those developed for simple, single-celled systems. Here, we engineered a set of gene regulatory tools by combining self-cleaving ribozymes with various upstream competing sequences that were designed to disrupt ribozyme self-cleavage. As a proof-of-concept, we were able to modulate GFP expression in mammalian cells, and then showed the feasibility of these tools in *Drosophila* embryos. For each system, the fold-reduction of gene expression was influenced by the location of the self-cleaving ribozyme/upstream competing sequence (i.e. 5’ untranslated region (UTR) vs. 3’UTR) and the competing sequence used. Together, this work provides a set of genetic tools that can be used to tune gene expression across various eukaryotic systems.

## INTRODUCTION

Synthetic biology is an interdisciplinary field that relies on biologists, engineers, mathematicians, and others to create novel biological systems by engineering and interchanging genetic parts derived from nature (1,2). This has led to advancements of various fields in medicine, molecular biology, and biotechnology. The ability to construct and analyze these systems has increased due to the availability of gene regulatory tools. Previous work has shown that these tools have the ability to regulate different steps of gene expression, including transcription (3), mRNA processing and stability (4), translation (5), and protein synthesis/stability (6). This ability has been particularly useful in the construction of synthetic gene circuits, such as counting devices (7), patterning devices (8), toggle switches (9), and gene oscillators (10), as well as the production of novel drugs, therapeutics, and biofuels.

While gene regulatory tools have been developed for various model systems, the development of these tools in eukaryotic systems has been outpaced compared to those developed in single-celled systems like bacteria and yeast. Initially, the development of gene regulatory tools in eukaryotic systems had been focused on transcriptional control (1). The tools to regulate transcription include the use of naturally-occurring (e.g. LacI, TetR, Gal4) and synthetic (e.g. zinc fingers, transcription activator-like effectors) transcription factors that have the ability to activate or inhibit gene expression (11–16). Later, other methods of gene regulation have been developed to control translation (upstream open reading frames (uORFs), microRNAs, aptamers) and protein turnover (17–23). More recently, clustered regularly interspaced short palindromic repeats (CRISPR) nucleases have been repurposed to act as synthetic transcription factors that have the ability to target virtually any gene of interest (24,25). Even with these tools available, more powerful tools are needed to precisely control gene expression within eukaryotic systems.

One promising gene regulatory tool that has the potential to fine-tune gene expression are self-cleaving ribozymes, which are natural RNA structures that are able to catalyze their own cleavage (26). When inserted into a transcript, these ribozymes reduce protein levels through self-cleavage and subsequent RNA degradation (Figure 1). Previous work has shown that inserting ribozymes in various loci of an mRNA transcript disrupts mRNA stability within bacteria, yeast, and mammalian cells (4,27,28). Previous work in bacteria has also shown that the insertion of sequences flanking a ribozyme and ribosome binding site can alter the ribozyme’s cleavage activity (29). Here, we used Mfold to engineer a set of genetic tools based on self-cleaving ribozymes that can be used to regulate gene expression in eukaryotic systems. By combining ribozymes with upstream competing sequences that have the potential to base-pair with a major stem of the ribozyme and prevent ribozyme self-cleavage (Figure 1B), we show that gene expression can be tuned in two model systems. We initially show that these tools can tune expression of a fluorescent reporter in HEK293T cells, and then we implemented the ribozyme constructs in *Drosophila* embryos. While we observed that these tools were able to modulate gene expression in two model systems, there was a lack of correlation between RNA secondary structure prediction algorithms and the experimental data. Together, these results show that self-cleaving ribozymes combined with upstream competing sequences can modulate gene expression in eukaryotic systems, and that other factors, besides ribozyme self-cleavage and base-pair interactions, influence gene expression.

**Figure 1:**
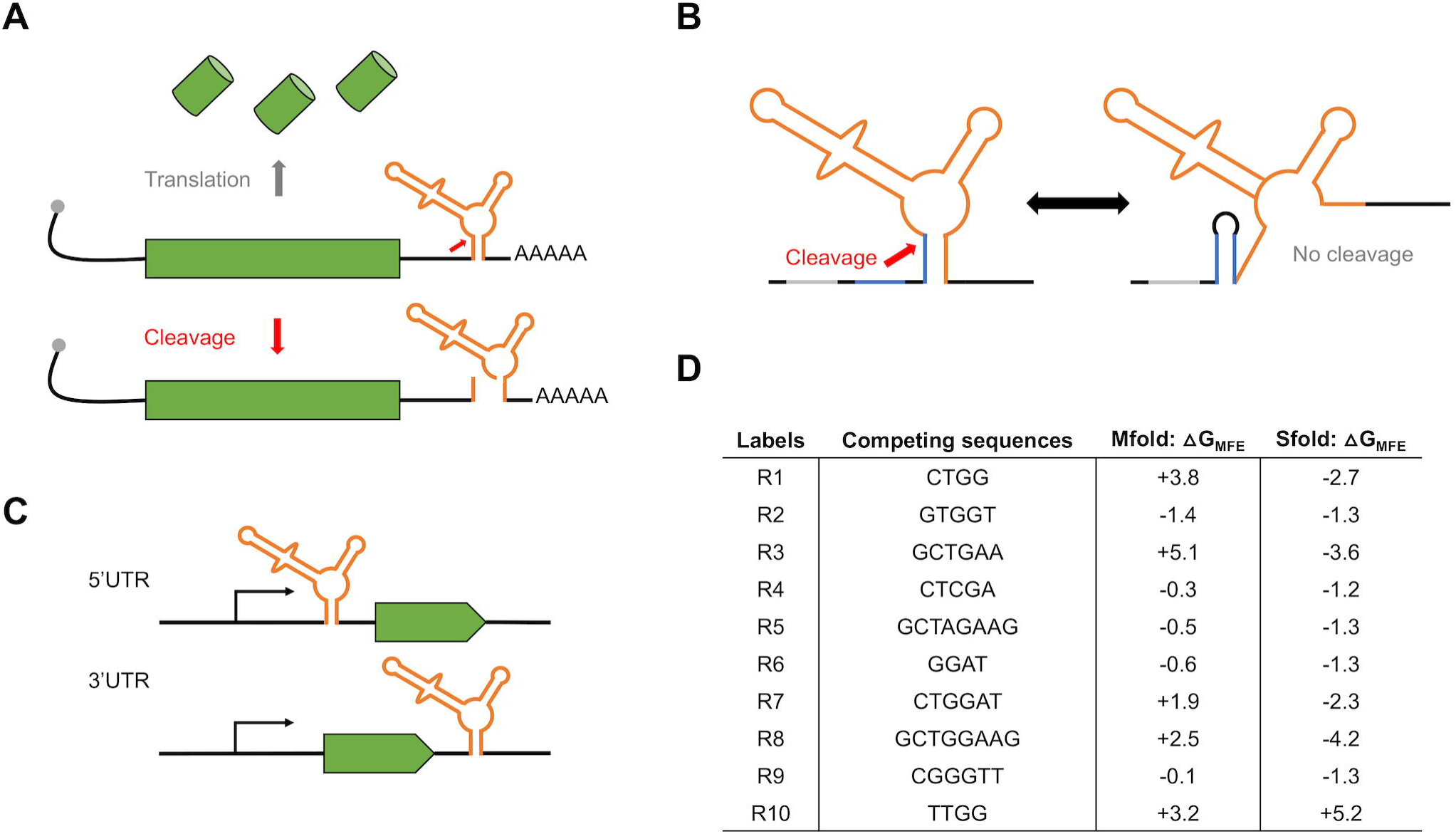
Gene regulatory tools based on self-cleaving ribozymes. **(A)** Inserting self-cleaving ribozymes in the 3’ untranslated region (UTR) of a gene leads to cleavage (red arrow) and subsequent mRNA transcript destabilization/decay and inhibition of protein synthesis. **(B)** Conceptual design of tunable self-cleaving ribozymes. A competing sequence (blue) is placed directly upstream of the ribozyme (orange). Base-pairing of the competing sequence with a part of the ribozyme stem prevents ribozyme self-cleavage. The ribozyme is flanked by insulating sequences (gray) to aid in preventing base-pairing interactions between the ribozyme and other sequences in the 3’UTR. **(C)** Schematic of the constructs used to test the ribozyme constructs in mammalian cells and *Drosophila*. We placed the ribozyme (orange) either in the 5’UTR or 3’UTR of the reporter genes used (green). **(D)** List of the competing sequences used in this study, along with their labels used in Figures 2 and 3. Also listed are the free energy differences between the minimal free energy structures of ribozymes in a cleavable and non-cleavable conformation for each competing sequence derived Mfold and Sfold. Note that R0 indicates a self-cleaving ribozyme lacking competing sequence.

**Figure 2:**
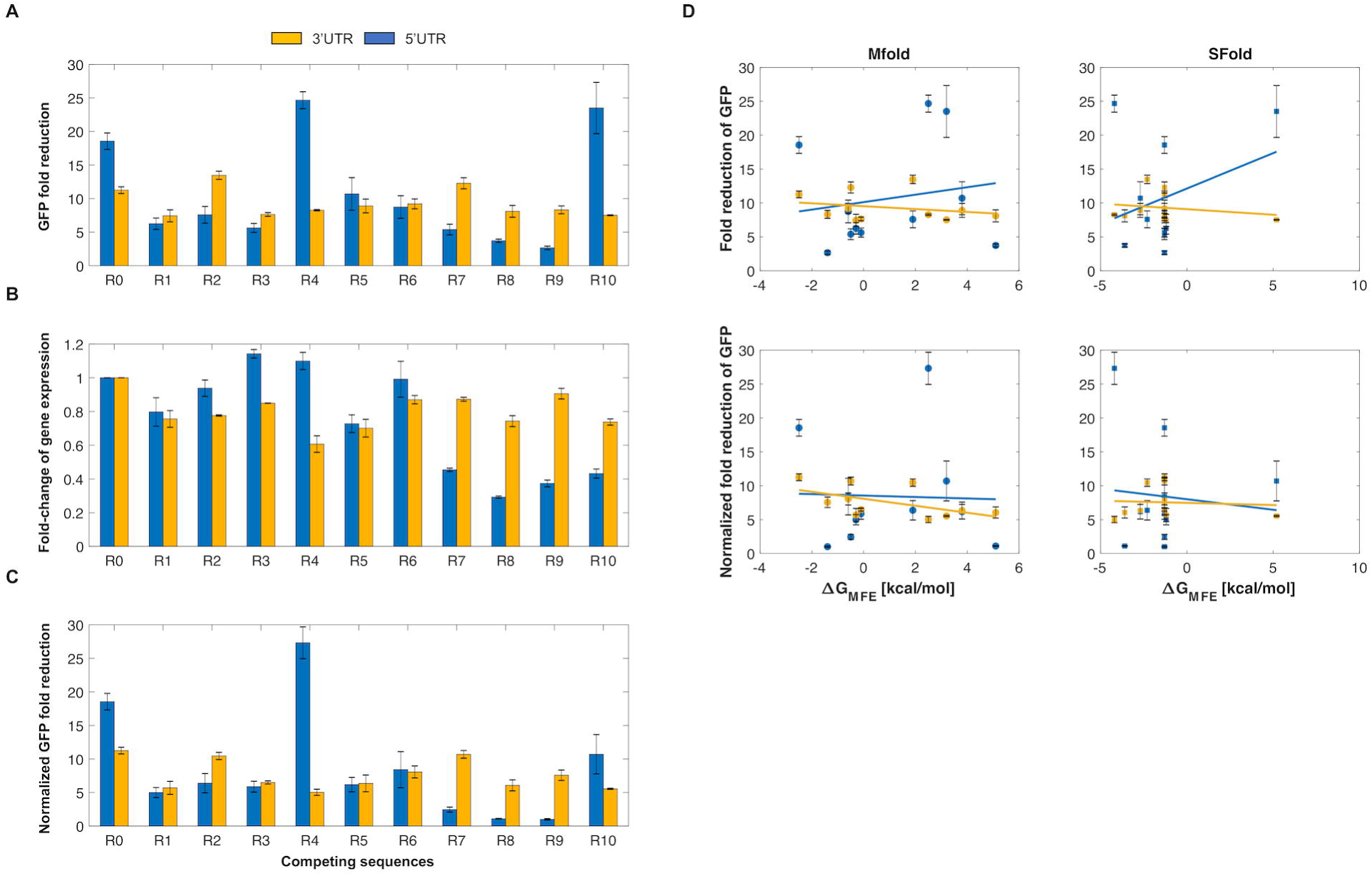
Self-cleaving ribozymes can tune gene expression in mammalian cells. **(A)** The average fold-reduction of GFP observed from the flow cytometry analysis for various competing sequences used in the 3’UTR (yellow) and 5’UTR (blue). The constructs were transiently transfected and incubated for 48 hours at 37°C. After incubation, the cells were trypsinized and resuspended in 1xPBS for flow cytometry analysis. **(B)** Comparison of the fold-change of GFP expression when a competing sequence is inserted in the 3’UTR (yellow) or 5’UTR (blue) of the transcript. A value of one indicates no change. **(C)** Normalized average of GFP fold-reduction using the data from Figures 2A/B. This represents the loss of reporter gene expression only due to ribozyme activity. All error bars represent the standard deviation from at least three independent transfections. Note that R0 indicates a self-cleaving ribozyme lacking competing sequence. **(D)** Predicted relationship between the fold-reduction of GFP and the free energy difference between cleavable and non-cleavable ribozyme conformations. Plots in column one and two compare the fold-reduction levels with the free energies calculated from Mfold and Sfold, respectively. The first and second rows represent the fold-reduction data (Figure 2A) and the normalized fold-reduction data (Figure 2C), respectively.

**Figure 3:**
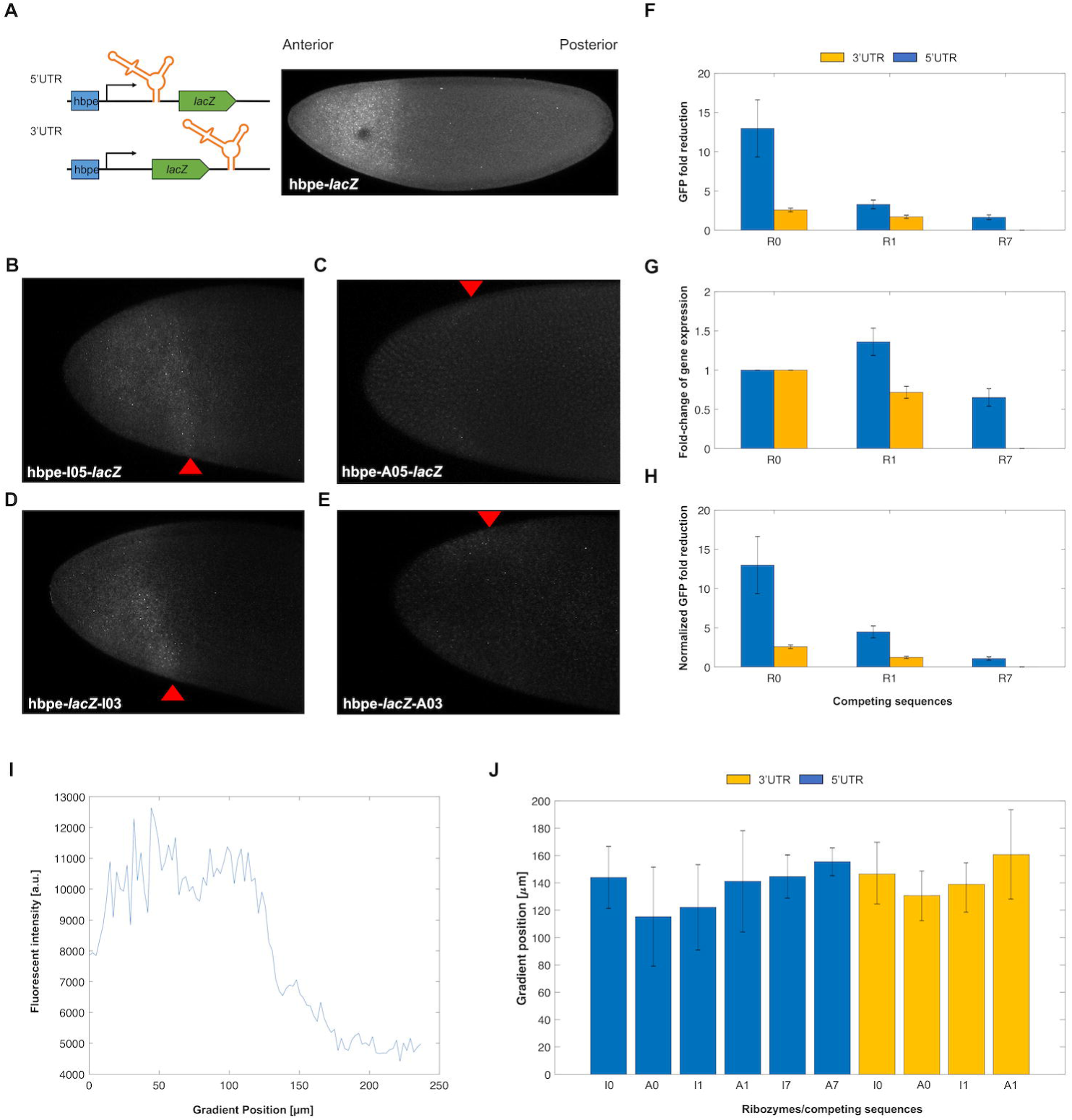
Self-cleaving ribozymes can tune gene expression in *Drosophila* embryos. **(A)** Depiction of the ribozyme constructs and its expression domain in *Drosophila* embryos. The domain of *lacZ* is similar to the endogenous *hunchback* (*hb*) gradient due to the hunchback proximal enhancer (hbpe). During early development, *hb* is strongly expressed in the anterior of the embryo. **(B-E)** Representative images of *in situ* hybridized *Drosophila* embryos probed for *lacZ*. Each embryo imaged expresses *lacZ* under the control of the hbpe and contains an inactive **(B/D)** or active **(C/E)** ribozyme. Red triangles represent the width of the *lacZ* gradient. **(F)** The average fold-reduction of *lacZ* observed from the imaging data for various competing sequences. Embryos were collected from transgenic fly lines constitutively expressing *lacZ from* the hbpe containing a ribozyme sequence in the 3’ UTR (yellow) or 5’ UTR (blue) and prepared for image analysis. Images were acquired using a Zeiss LSM710 confocal microscope. **(G)** Fold-change of *lacZ* expression due to effects other than ribozyme activity. A value of 1 indicates no change. **(H)** Normalized average fold-reduction of *lacZ* using the data from Figures 3F/G. This represents the reduction of *lacZ* expression solely due to ribozyme activity. All error bars represent the standard deviation from at least 10 embryos. Note that R0 indicates a self-cleaving ribozyme lacking an upstream competing sequence. Also note that fly lines containing the R7 competing sequence in the 3’UTR were not analyzed. **(I)** The fluorescence intensity at various positions of the embryo. A domain width of zero indicates the anterior pole and increasing values indicate a position closer to the posterior. **(J)** The average width of the *lacZ* domain for each ribozyme and competing sequence listed.

## MATERIALS AND METHODS

### Strains, plasmids, oligonucleotides, and fly lines

All strains, plasmids, oligos, gBlocks, and fly lines used in this work can be found in **Supplementary Document 1**. All PCR amplifications were performed using Q5 Hot Start High-Fidelity 2X Master Mix (NEB, Cat: M0494S) unless specified. All fly lines were generated using site-specific PhiC31-mediated insertion from Genetivision.

We used the pcDNA3.1^(+)^ mammalian expression vector (Thermo Fisher, Cat: V79020) for expression of GFP in HEK293T cells. For this study we used the hammerhead self-cleaving ribozyme from *Schistosoma mansoni* as it has been associated with high catalytic activity *in vitro* and *in vivo* (29,30). We first built the active ribozyme constructs by first PCR amplifying GFP and inserting it into the NotI and PstI sites of pCB1180. The inactive ribozyme constructs were built by creating a single point mutation that abolishes catalytic activity of the ribozyme (31). Then, annealed and phosphorylated oligos containing the inactive and active ribozymes were inserted into the XhoI and NotI sites, located in the 5’ untranslated region (UTR), to make pCB1134/1135. To insert these ribozyme-GFP sequences into pcDNA3.1^(+)^, we PCR amplified the ribozyme-GFP sequence from pCB1134/1135 and inserted it into the HindIII and XbaI restriction sites in pcDNA3.1^(+)^ to create pCB1136/1137. The upstream competing sequences were inserted into pCB1136/1137 by linearizing the plasmids with EcoRI and XhoI and then ligating with annealed and phosphorylated oligos containing the competing sequences of interest (Figure 1D). For insertion of the ribozyme/upstream competing sequences in the 3’UTR of GFP, the ribozyme/upstream competing sequences were PCR amplified from the previously built 5’UTR constructs and inserted into the XbaI site of pCB1133.

We used the pUAST-attB *Drosophila* expression vector (Drosophila Genomics Resource Center, Cat: 1419) for creating the transgenic fly lines containing the *lacZ* reporter. To generate the ribozyme constructs, we first removed the UAS-hsp70 sequence using the HindIII and KpnI restriction sites and added the hunchback (hb) proximal enhancer (hbpe), the eve minimal promoter, and the *lacZ* reporter to create pCB1181. Expressing *lacZ* from the hbpe creates a well-established domain of *hb* to easily study the effects from the self-cleaving ribozymes (32–34). For the insertion of the self-cleaving ribozymes into the 5’UTR and 3’UTR of *lacZ*, the StuI and KpnI restriction sites of pCB1181 were used, respectively. To insert the upstream competing sequences, both the EcoRI and AvrII sites were added upstream of the ribozyme sequence for ligation with annealed and phosphorylated oligos containing the competing sequences of interest.

### Predicting secondary structures of self-cleaving ribozymes/upstream competing sequences

The online tools Mfold and Sfold were used to predict the minimal free energy (MFE) structures of the ribozymes lacking or containing an upstream competing sequence using the default settings (35,36). We extracted the ΔG of the structures associated with the lowest free energy of a ribozyme in a cleavable and non-cleavable conformation. The ΔG of each upstream competing sequence was calculated as the difference between the ΔG of the cleaved and non-cleaved structures. See **Supplementary Figure 1** for a representative secondary structure of ribozymes in a cleaving or non-cleaving conformation.

### Transient transfections of pcDNA3.1^(+)^-ribozyme constructs

Transfection-grade DNA was prepared using the QIAGEN Plasmid Mini Kit (QIAGEN, Cat: 12125). One day prior to the transient transfections, HEK239T cells were seeded onto either 35mm or 24-well plates with complete media (Dulbecco’s Modified Eagle Medium (Invitrogen, Cat: 11965-092) supplemented with 10% fetal bovine serum (Invitrogen, Cat: A3840001)). Each pcDNA3.1^(+)^-ribozyme construct was transiently transfected using FuGeneHD (Promega, Cat: E2311). Cells were then incubated for 48 hours prior to preparing the cells for flow cytometry. See **Supplementary Table 1** for details of the transient transfections performed using each plate format.

### Flow cytometry analysis of transiently transfected HEK293T cells

We trypsinized the transiently transfected HEK293T cells using trypsin-EDTA (Thermo Fisher, Cat: 25200056) and resuspended them in 500mL 1xPBS (Fisher Scientific, Cat: MT21040CV). The cells were analyzed for fluorescence using the Accuri C6 Flow Cytometer with CFlow plate sampler (Becton Dickinson). The events were gated based on the forward scatter and side scatter, with fluorescence measured in FL2-H, using the 533/30 filter, from at least 10,000 gated events. The fold-reduction of GFP was calculated as the ratio of the fluorescence values for the cells transfected with an inactive ribozyme with a specific competing sequence over that of an active ribozyme with the same competing sequence.

### Fluorescent *in situ* hybridization (FISH) of *Drosophila* embryos

All embryos were aged to 2-4 hours from laying and then fixed using 37% formaldehyde following standard protocols (37). FISH was combined with fluorescent immunostaining following standard protocols (37). Briefly, fixed embryos were washed in 1xPBS buffer supplemented with 0.05% Tween-20, and then hybridized with a fluorescein (ftc)-conjugated anti-sense *lacZ* probe at 55°C. The embryos were washed and incubated with the rabbit anti-histone (Abcam, Cat: ab1791) (1:10,000 dilution) and goat anti-ftc (Rockland, Cat: 600-101-096) (1:5,000 dilution) primary antibodies overnight at 4°C. Embryos were then washed and incubated for 1.5 hours with fluorescent donkey anti-rabbit-546 (Invitrogen, Cat: A10040) (1:500 dilution) and donkey anti-goat-647 (Invitrogen, Cat: A21447) (1:500 dilution) secondary antibodies at room temperature. Finally, the embryos were washed and stored in 70% glycerol at −20°C prior to being imaged. All prepared embryos were imaged within two weeks of protocol completion.

### Imaging and image analyses of *Drosophila* embryos

To reduce variability from the fluorescence measurements, the intensity output of the 488 nM laser was used for laser calibration prior to embryo imaging (38). The calibration was performed by measuring the intensity of the 488 nM laser through the transmitted light channel giving us the output strength of the laser. This allowed us to compensate for potential variability of laser strength between imaging sessions. The prepared embryos were mounted laterally using 70% glycerol using two pieces of double-sided tape. A Zeiss LSM 710 microscope was used to acquire 15-25 z-slices 45-60 μm apart at 40x magnification.

Using Fiji, the z-max intensity projection for each embryo was measured for its fluorescence intensity. The *hb* expression domain was used as the cutoff for signal, as the expression profile of *lacZ* should match the endogenous *hb* expression pattern due to expression from its enhancer (hbpe). The fluorescent signal was obtained by measuring the intensity from the anterior pole to the edge of *hb* domain using the tools available in Fiji. After measuring signal, background noise was measured as the intensity outside of the *hb* expression pattern. The fold-reduction of *lacZ* was calculated as the ratio of the fluorescence values for the embryos with an inactive ribozyme with a specific competing sequence over that of an active ribozyme with the same competing sequence. Refer to **Supplementary Document 2** for an in-depth protocol.

Using the same embryos, the width of the *lacZ* gradient was compared with the active and inactive ribozyme constructs. For this analysis, we used a supervised MATLAB script to first locate and orient the embryo, and then shape the embryos’ periphery boundary. We then measured the fluorescence of the embryo across the anterior-posterior axis (see supplementary material for MATLAB scripts). To measure the distance from the anterior pole to the boundary of the *lacZ* domain, we selected three points along the y-axis and extracted the width corresponding to 50% loss of the maximum intensity. We selected three different y-values to account for asymmetrical *lacZ* gradients (**Supplementary Figure 3**). The median of the three values was used to represent the measurement of the *lacZ* gradient.

## RESULTS

### Designing self-cleaving ribozymes containing tunable upstream competing sequences

For this study, we used the hammerhead self-cleaving ribozyme as it has shown high activity *in vitro* and *in vivo* (29,30). Though these ribozyme constructs can be placed in various locations within a transcript, we chose to test two specific locations: the 5’ and 3’UTR of the reporter genes tested (Figure 1C). The competing sequences were placed upstream of the ribozyme to ensure that transcription of the ribozyme before the competing sequence did not result in self-cleavage prior to the transcription of the competing sequence. Insulating sequences were flanked upstream of the ribozyme/competing sequence to limit ribozyme misfolding due to flanking sequences (Figure 1B). Finally, we designed the competing sequences using Mfold (35) to obtain a set of sequences that were associated with varying levels of predicted folded and misfolded ribozyme structures (Figure 1D). Each competing sequence varied in sequence length and composition and were associated with different propensities to base-pair with the stem of the ribozyme. Finally, each competing sequence lacked a start codon to prevent premature translation initiation.

### Self-cleaving ribozymes combined with upstream competing sequences can modulate gene expression in mammalian cells

We first sought to test these ribozyme constructs in a mammalian system. To this end, we tested the ribozyme constructs in HEK293T cells. We inserted the self-cleaving ribozymes and 10 different upstream competing sequences in the 5’ UTR or the 3’UTR of GFP to observe how various sequence configurations impacted reporter gene expression (Figure 2). For each ribozyme/competing sequence tested, we used an inactive ribozyme with the same competing sequence to act as a control. As the inactive and active ribozymes only differ by a single point mutation (31), the overall structure of the ribozyme was preserved. After transiently transfecting these reporter constructs, the fluorescence of the cells was analyzed by flow cytometry analysis. We found that these ribozyme constructs were able to reduce expression of GFP in HEK293T cells, with fluorescence generally being associated in a bimodal distribution (untransfected cells and cells associated with varying GFP levels) (**Supplementary Figure 2**). When located in the 5’UTR, the ribozymes/upstream competing sequences generally resulted in greater range of fold-reduction levels compared to when located in the 3’UTR (Figure 2A).

As the GFP fold-reduction levels between the 5’ and 3’UTR constructs were variable, we wanted to assess the effect of competing sequence insertion on GFP expression. Due to prior work showing that the formation of secondary structures strongly effects transcript stability (39), we compared the fluorescence of the cells transiently transfected with ribozyme constructs containing an inactive ribozyme lacking an upstream competing sequence to that of inactive ribozymes containing an upstream competing sequence (Figure 2B). While the loss of GFP expression was fairly consistent for the constructs containing ribozymes with competing sequences in the 3’UTR (~20-40% loss of GFP expression), GFP expression loss was more noticeable when the ribozyme/competing sequences were placed in the 5’UTR. When placed in the 5’UTR, the loss of gene expression ranged from negligible loss (e.g. R2, R6) to ~70% loss (e.g. R8) (Figure 2B). Interestingly, the insertion of some upstream competing sequences resulted in increased expression of GFP (e.g. R3, R4). We then accounted for the loss of gene expression due to the insertion of a competing sequence by normalizing the fold-reduction data from Figure 2A using the data from Figure 2B (Figure 2C). While this generally resulted in less fold-reduction of each construct, a wide dynamic range was generally maintained, from almost no fold-reduction to ~25-fold-reduction of GFP.

After obtaining the experimental data, we then sought to gain insight into the relationship between the fold-reduction of gene expression and the predicted energies of misfolding. To this end, we compared the GFP fold-reduction levels with the predicted free energy differences obtained from Mfold. To obtain these values, the difference between the ΔG associated with the MFE structure of a ribozyme in a cleavable conformation and the ΔG associated with the MFE in a non-cleavable conformation was calculated (**Supplementary Figure 1**). While the experimental data from HEK293T cells showed a wide dynamic range of fold-reduction levels, there was a lack of correlation between the experimental data and predicted free energy differences (Figure 2D). We then sought to use a different RNA predictive folding algorithm to see if it could better correlate the fold-reduction of gene expression to predicted free energies. Thus, we used Sfold to compare MFE’s to the GFP fold-reduction (36). Similar to Mfold, there was a lack of correlation between the experimental fold-reductions to the predicted free energies (Figure 2D). The lack of a correlation indicates the presence of external factors that influence ribozyme self-cleavage, thus currently making this approach non-predictive. Even so, our data show that ribozyme/upstream competing sequences can be used to tune gene expression in mammalian cells.

### Self-cleaving ribozymes/upstream competing sequences can modulate gene expression in *Drosophila*

As a proof-of-concept, we next wanted to test these tools in a multicellular system. We chose to work with *Drosophila* embryos as we have previously used this system to study synthetic networks (40). Thus, we generated transgenic fly lines carrying these ribozyme constructs. We first designed *Drosophila* expression vectors containing the *lacZ* reporter expressed from the hunchback proximal enhancer (hbpe). This enhancer results in an expression pattern similar to endogenous *hb*, which has a sharp boundary at roughly 50% anterior-to-posterior (AP) coordinate (32–34). The hbpe drives expression with a boundary at roughly 33% AP coordinate (Figure 3A), which allowed us to quantitatively test these ribozymes *in vivo*. Similar to the work in HEK293T cells, each ribozyme/upstream competing sequence tested were compared to an inactive ribozyme containing the same competing sequence to act as a negative control. Embryos were first hybridized with an antisense *lacZ* probe, then imaged by confocal microscopy. We found that the insertion of ribozyme/competing sequences into a transcript expressing *lacZ* were able to tune *lacZ* expression levels in *Drosophila* embryos (Figures 3B-E). Unlike with the mammalian cell data, normalizing the fold-reduction data by accounting for the effects of inserting the upstream competing sequences on *lacZ* expression resulted in negligible changes to the measured dynamic range of fold-reduction values (Figures 2A-C, 3F-G). While the fold-reduction values observed in *Drosophila* were generally less than those observed in HEK293T cells, the correlation of fold-reduction values, compared to the work in mammalian cells, remained the same (i.e. R0 > R1 > R7) and maintained a high dynamic range (~2-14 fold-reduction of *lacZ*).

Using the same images, we then compared the width of the *lacZ* domain along the anterior-posterior axis. We hypothesized that the embryos containing an active ribozyme construct would be associated with a reduced domain width as the expression of *lacZ* would be reduced at locations containing weak fluorescent intensity (i.e. distal to anterior pole). For each ribozyme construct, we observed that the differences in the *lacZ* domain width were small, but noticeable across all constructs. Interestingly, only the two strongest ribozymes (i.e. A0-5UTR, A0-3UTR) resulted in a noticeable *lacZ* gradient reduction (Figure 3J), though the average gradient width between active and inactive ribozymes were not statistically different. These results indicate that the *lacZ* domain width did not vary between active and inactive ribozymes regardless of location or competing sequence.

## DISCUSSION

In this work, we engineered a set of genetic tools that were able to modulate gene expression in HEK293T cells and *Drosophila* embryos. At face value, inserting the ribozymes in the 5’UTR of the reporter genes yielded a greater range of fold-reduction levels compared to the 3’UTR. However, we observed that insertion of upstream competing sequences resulted in the inhibition of gene expression in the absence of ribozyme self-cleavage. This effect was greater when the ribozyme/competing sequence was located in the 5’UTR (Figure 2B). After normalizing the fold-reduction levels by accounting for the loss of gene expression, we observed that some ribozyme constructs (most notably the 5’UTR constructs) reduced gene expression more weakly compared to that data prior to normalization (Figures 2A/C). In general, the ribozymes/upstream competing sequences were observed to reduce gene expression more strongly in HEK293T cells compared to *Drosophila* embryos (Figures 2 and 3), which has also been observed in recent work (41). This difference could be due to different biological machinery between mammalian and insect models, different experimental assays, or the constructs themselves, as they contain different promoters and reporter genes. Even with the differences in fold-reduction levels between these model systems, these tools maintained a dynamic range of gene expression regulation (~1-25 in HEK293T cells and ~2-14 in *Drosophila*). While the experimental data did not show a high correlation with the RNA secondary structure predictions (Figure 2D), we provide a set of gene regulatory tools based on empirical measurements between competing sequences and strength of gene reduction.

Prior to experimental work, we used Mfold (35,36) to design a set of competing sequences that were associated with a wide range of free energies (Figure 1D). When comparing these predicted free energies to the fold-reduction levels observed in our experimental data (Figures 2A/C), we generally observed a weak correlation (Figure 2D). This discrepancy could have been due to a variety of factors. For instance, the insulating sequences, used to prevent interactions between the ribozyme and flanking sequences, could have affected the ability of the competing sequences to base-pair with the ribozyme. While Mfold and Sfold predictions showed minimal interactions between the ribozyme and insulating sequences, the sequences flanking the insulating sequences could have interacted with the competing sequence, ribozyme, and/or the insulating sequence. It is also possible that one or more of the competing and/or insulating sequences contain a target sequence for a native biological factor or pathway, such as an endogenous transcription factor, internal ribosome entry site (IRES), or RNAi. While the addition of a specific target sequence is unlikely, novel transcription factors, IRES’, and non-coding RNAs continue to be discovered in eukaryotic systems, including *Drosophila* (42–48). Finally, Mfold and/or Sfold may lack the ability to predict the fold-reduction of gene expression associated with the ribozyme constructs. Recent work has shown that hammerhead ribozymes are associated with varying cleavage activities across different model systems (e.g. mammalian vs. yeast) and experimental setups (e.g. *in vitro* vs. *in vivo*) (41), which show that cellular context is likely important for the observed activity. Another possibility is that Mfold and Sfold are not accurately capturing RNA folding. While algorithms, such as Mfold and Sfold, have the ability to predict RNA secondary structures, ribozymes can form complex 3D structures (e.g. pseudoknots) that cannot be predicted accurately. Due to these reasons, current predictive RNA folding algorithms may not be sufficient for accurate secondary structure predictions. Improvements on RNA structure prediction models will allow for a more accurate design of competing sequences.

Experimental data indicated that insertion of the upstream competing sequences generally inhibited gene expression when compared with the constructs lacking these sequences. This phenomenon could be due to various reasons. For one, the mRNA transcripts could have been subjected to the no-go decay pathway (49). This mRNA surveillance pathway occurs when ribosomes have stalled during translation, resulting in cleavage and subsequent degradation. While some of the ribozyme constructs resulted in drastic reduction of gene expression from competing sequence insertion, the majority of these constructs had a small, but noticeable effect on gene expression (Figure 2B). Similar to the RNA folding algorithms, another possibility could be that one or more of the competing sequences was a target sequence for an endogenous biological factor or pathway. To prevent variation of gene expression when using these ribozyme constructs, longer insulating sequences can be flanked to both the 5’ and 3’ ends of the ribozyme/upstream competing sequences. This could prevent interactions between the ribozyme or competing sequence with flanking sequences, resulting in fold-reduction levels only from ribozyme self-cleavage.

## CONCLUSIONS

We developed a set of tools that were able to tune gene expression in HEK293T cells and *Drosophila*. While the free energies obtained from the predictive RNA secondary structure tool did not correlate well with the experimental data, the competing sequences used in this work provide a set of genetic tools associated with a wide range of fold-reduction levels. Though tested in mammalian and insect systems, these tools should be applicable in other eukaryotic systems, such as *C. elegans*, zebrafish, and mice. Previous work has shown that self-cleaving ribozymes are found naturally in these organisms (50–52) and have been used for therapeutic applications (4,53). These tools will be useful for studies involving synthetic biology, especially for the purposes of building and studying synthetic gene circuits.

## ACKNOWLEDGMENTS

The pUAST-attB plasmid was a gift from the Drosophila Genomics Resource Center, who are funded from the National Institutes of Health (2P40OD010949). This work was supported by the U.S. Department of Education [Graduate Assistance in Areas of National Need Biotechnology Fellowship (P200A140020)] and the National Science Foundation (MCB-1413044).

## AUTHOR CONTRIBUTIONS

AAJ, TJ, CLB, and GTR conceived and planned the experiments. TJ and GY carried out the molecular cloning. TJ performed the transient transfections and flow cytometry. TJ and GY carried out the *in situ* hybridizations. HA imaged the embryos. TJ performed the data analysis. TJ, GTR, and CB wrote the manuscript with input from all authors.

## SUPPORTING INFORMATION

**Supplementary Figure 1:** Representative figures depicting secondary structures of self-cleaving ribozymes. **(A)** Self-cleaving ribozyme that lacks a competing sequence in a cleavable conformation. **(B)** Self-cleaving ribozyme that contains a competing sequence in a cleavable conformation. **(C)** Self-cleaving ribozyme that contains a competing sequence in a non-cleavable conformation. The red text indicates the insulating sequence, green text indicates the competing sequence, and black text indicates the ribozyme.

**Supplementary Figure 2:** Flow cytometry data of transiently transfected HEK293T cells. **(A)** Representative forward and side scatter plot of HEK293T cells transiently transfected with ribozyme constructs. The cell population was gated in green. **(B)** Histograms of transiently transfected HEK293T cells. Plotted are the number of cells at corresponding fluorescent values of untransfected cells (black), cells containing an active ribozyme/competing sequence (red), and cells containing an inactive ribozyme/competing sequence (blue) in the 5’UTR (top row) or 3’UTR (bottom row) of *gfp*.

**Supplementary Figure 3:** Representative embryos labeled with *lacZ* gradient width associated with **(A/B)** symmetric and **(C/D)** asymmetric *lacZ* gradients. Red line indicates end of *lacZ* gradient. Multiple red lines indicate the width of the lacZ gradient at a particular anterior-posterior axis length.

**Supplementary Table 1:** Transfection conditions used for 35mm plates and 24-well plates.

**Supplementary Document 1:** DNA constructs and fly lines used in this work.

**Supplementary Document 2:** In-depth protocol for measuring fluorescence intensity of *Drosophila* embryos.

## REFERENCES

1. Lienert F, Lohmueller JJ, Garg A, Silver PA. Synthetic biology in mammalian cells: Next generation research tools and therapeutics. Nat Rev Mol Cell Biol. 2014;15:95–107.

2. Singh V. Recent advancements in synthetic biology: Current status and challenges. Gene. 2014;535:1–11.

3. Guzman L, Belin D, Carson MJ, Beckwith J. Tight regulation, modulation, and high-level expression by vectors containing the arabinose PBAD promoter. J Biotechnol. 1995;177:4121–30.

4. Yen L, Svendsen J, Lee JS, Gray JT, Magnier M, Baba T, et al. Exogenous control of mammalian gene expression through modulation of RNA self-cleavage. Nature. 2004;431:471–6.

5. Salis HM, Mirsky EA, Voigt CA. Automated design of synthetic ribosome binding sites to control protein expression. Nat Biotechnol. 2009;27:946–50.

6. McGinness KE, Baker TA, Sauer RT. Engineering controllable protein degradation. Mol Cell. 2006;22:701–7.

7. Friedland AE, Lu TK, Wang X, Shi D, Church G, Collins JJ. Synthetic gene networks that count. Science. 2009;324:1199–202.

8. Basu S, Gerchman Y, Collins CH, Arnold FH, Weiss R. A synthetic multicellular system for programmed pattern formation. Nature. 2005;434:1130–4.

9. Gardner TS, Cantor CR, Collins JJ. Construction of a genetic toggle switch in *Escherichia coli*. Nature. 2000;403:339–42.

10. Elowitz MB, Leibler S. A synthetic oscillatory network of transcriptional regulators. Nature. 2000;403:335–338.

11. Brown M, Figge J, Hansen U, Wright C, Jeang K-T, Khoury G, et al. Lac repressor can regulate expression from a hybrid SV40 early promoter containing a lac operator in animal cells. Cell. 1987;49:603–12.

12. Gossen M, Bujard H. Tight control of gene expression in mammalian cells by tetracycline-responsive promoters. Proc Natl Acad Sci. 1992;89:5547–51.

13. Maeder ML, Thibodeau-Beganny S, Osiak A, Wright DA, Anthony RM, Eichtinger M, et al. Rapid “open-source” engineering of customized zinc-finger nucleases for highly efficient gene modification. Mol Cell. 2008;31:294–301.

14. Morbitzer R, Römer P, Boch J, Lahaye T. Regulation of selected genome loci using de novo-engineered transcription activator-like effector (TALE)-type transcription factors. Proc Natl Acad Sci. 2010;107:21617–22.

15. Garg A, Lohmueller JJ, Silver PA, Armel TZ. Engineering synthetic TAL effectors with orthogonal target sites. Nucleic Acids Res. 2012;40:7584–95.

16. Kakidani H, Ptashne M. GAL4 activates gene expression in mammalian cells. Cell. 1988;52:161–7.

17. Medenbach J, Seiler M, Hentze MW. Translational control via protein-regulated upstream open reading frames. Cell. 2011;145:902–13.

18. Ferreira JP, Overton KW, Wang CL. Tuning gene expression with synthetic upstream open reading frames. Proc Natl Acad Sci. 2013;110:11284–9.

19. Xie Z, Wroblewska L, Prochazka L, Weiss R, Benenson Y. Multi-input RNAi-based logic circuit for identification of specific cancer cells. Science. 2011;333:1307–11.

20. Ausländer S, Ausländer D, Müller M, Wieland M, Fussenegger M. Programmable single-cell mammalian biocomputers. Nature. 2012;487:123–7.

21. Nishimura K, Fukagawa T, Takisawa H, Kakimoto T, Kanemaki M. An auxin-based degron system for the rapid depletion of proteins in nonplant cells. Nat Methods. 2009;6:917–22.

22. Bonger KM, Chen L, Liu CW, Wandless TJ. Small-molecule displacement of a cryptic degron causes conditional protein degradation. Nat Chem Biol. 2011;7:531–7.

23. Banaszynski LA, Chen L-C, Maynard-Smith LA, Ooi AGL, Wandless TJ. A rapid, reversible, and tunable method to regulate protein function in living cells using synthetic small molecules. Cell. 2006;126:995–1004.

24. Qi LS, Larson MH, Gilbert LA, Doudna JA, Weissman JS, Arkin AP, et al. Repurposing CRISPR as an RNA-guided platform for sequence-specific control of gene expression. Cell. 2013;152:1173–83.

25. Gilbert LA, Larson MH, Morsut L, Liu Z, Brar GA, Torres SE, et al. CRISPR-mediated modular RNA-guided regulation of transcription in eukaryotes. Cell. 2013;154:442–51.

26. Ferré-D’Amaré AR, Scott WG. Small self-cleaving ribozymes. Cold Spring Harb Perspect Biol. 2010;2:a003574.

27. Link KH, Guo L, Ames TD, Yen L, Mulligan RC, Breaker RR. Engineering high-speed allosteric hammerhead. Biol Chem. 2007;388:779–86.

28. Win MN, Smolke CD. A modular and extensible RNA-based gene-regulatory platform for engineering cellular function. Proc Natl Acad Sci. 2007;104:14283–8.

29. Carothers JM, Goler J a, Juminaga D, Keasling JD. Model-driven engineering of RNA devices to quantitatively program gene expression. Science. 2011;334:1716–9.

30. Ferbeyre G, Smith JM, Cedergren R. Schistosome satellite DNA encodes active hammerhead ribozymes. Mol Cell Biol. 1998;18:3880–8.

31. Seela F, Debelak H, Usman N, Burgin A, Beigelman L. 1-Deazaadenosine: synthesis and activity of base-modified hammerhead ribozymes. Nucleic Acids Res. 1998;26:1010–8.

32. Driever W, Nüsslein-Volhard C. The bicoid protein is a positive regulator of hunchback transcription in the early *Drosophila* embryo. Nature. 1989;337:138–43.

33. Lehmann R, Nüsslein-Volhard C. hunchback, a gene required for segmentation of an anterior and posterior region of the *Drosophila* embryo. Dev Biol. 1987;119:402–17.

34. Perry MW, Bothma JP, Luu RD, Levine M. Precision of hunchback expression in the *Drosophila* embryo. Curr Biol. 2012;22:2247–52.

35. Zuker M. Mfold web server for nucleic acid folding and hybridization prediction. Nucleic Acids Res. 2003;31:3406–15.

36. Ding Y, Chan CY, Lawrence CE. Sfold web server for statistical folding and rational design of nucleic acids. Nucleic Acids Res. 2004;32:135–41.

37. Kosman D, Mizutani CM, Lemons D, Cox WG, McGinnis W, Bier E. Multiplex detection of RNA expression in *Drosophila* embryos. Science. 2004;305:846.

38. Liberman LM, Reeves GT, Stathopoulos A. Quantitative imaging of the Dorsal nuclear gradient reveals limitations to threshold-dependent patterning in *Drosophila*. Proc Natl Acad Sci. 2009;106:22317–22322.

39. Carrier TA, Keasling JD. Controlling messenger RNA stability in bacteria: Strategies for engineering gene expression. Biotechnol Prog. 1997;13:699–708.

40. Jermusyk AA, Murphy NP, Reeves GT. Analyzing negative feedback using a synthetic gene network expressed in the *Drosophila* melanogaster embryo. BMC Syst Biol. 2016;10:85.

41. Wurmthaler LA, Klauser B, Hartig JS. Highly motif- and organism-dependent effects of naturally occurring hammerhead ribozyme sequences on gene expression. RNA Biol. 2018;15:231–41.

42. Young RS, Marques AC, Tibbit C, Haerty W, Bassett AR, Liu JL, et al. Identification and properties of 1,119 candidate LincRNA loci in the *Drosophila melanogaster* genome. Genome Biol Evol. 2012;4:427–42.

43. Inagaki S, Numata K, Kondo T, Tomita M, Yasuda K, Kanai A, et al. Identification and expression analysis of putative mRNA-like non-coding RNA in *Drosophila*. Genes to Cells. 2005;10:1163–73.

44. Tupy JL, Bailey AM, Dailey G, Evans-Holm M, Siebel CW, Misra S, et al. Identification of putative noncoding polyadenylated transcripts in *Drosophila melanogaster*. Proc Natl Acad Sci. 2005;102:5495–500.

45. Marr II MT, D’Alessio JA, Puig O, Tjian R. IRES-mediated functional coupling of transcription and translation amplifies insulin receptor feedback. Genes Dev. 2007;21:175–83.

46. Maier D, Nagel AC, Preiss A. Two isoforms of the Notch antagonist Hairless are produced by differential translation initiation. Proc Natl Acad Sci. 2002;99:15480–5.

47. Rhee DY, Cho D-Y, Zhai B, Slattery M, Ma L, Mintseris J, et al. Transcription factor networks in Drosophila melanogaster. Cell Rep. 2014;8:2031–43.

48. Wang C, Yeung F, Liu P-C, Attar RM, Geng J, Chung LWK, et al. Identification of a novel transcription factor, GAGATA-binding protein, involved in androgen-mediated expression of prostate-specific antigen. J Biol Chem. 2003;278:32423–30.

49. Doma MK, Parker R. Endonucleolytic cleavage of eukaryotic mRNAs with stalls in translation elongation. Nature. 2006;440:561–4.

50. Webb C-HT, Riccitelli NJ, Ruminski DJ, Lupták A. Widespread occurrence of self-cleaving ribozymes. Science. 2009;326:953.

51. Martick M, Horan LH, Noller HF, Scott WG. A discontinuous hammerhead ribozyme embedded in a mammalian messenger RNA. Nature. 2008;454:899–902.

52. Salehi-Ashtiani K, Szostak JW. In vitro evolution suggests multiple origins for the hammerhead ribozyme. Nature. 2001;414:82–4.

53. Peace BE, Florer JB, Witte D, Smicun Y, Toudjarska I, Wu G, et al. Endogenously expressed multimeric self-cleaving hammerhead ribozymes ablate mutant collagen *in cellulo*. Mol Ther. 2005;12:128–36.

