## Supplementary Document 2 for "Tunable self-cleaving ribozymes for modulating gene expression in eukaryotic systems"

**Protocol for measuring fluorescence intensity of embryos (associated with Figures 4F-H)**

1. Download, install, and open Fiji program (<https://imagej.net/Fiji/Downloads>)
2. Import the image into Fiji by dragging the image file to the opened window.
3. Select Image > Stacks > Z Project…
4. The default “Start slice” and “Stop slice” is fine. It will default to the minimum and maximum number of slices, respectively. Select “Projection type” to “Max Intensity” and click “Okay”. See Figure A for representative image.
5. Use the “Polygon selections” to draw an area of *hunchback* (*hb*) signal. See Figure B below for representative embryo bounded by *hb* domain (yellow).
6. Select Analyze > Measure and record the “Mean” value to obtain the average intensity of the *hb* domain.
7. Select Edit > Draw to keep the outline of the *hb* domain but allow for the addition of the background noise selection. See Figure C below for representative embryo bounded by background noise domain (yellow). The *hb* domain is drawn (red) to ensure signal is differentiated from background noise.

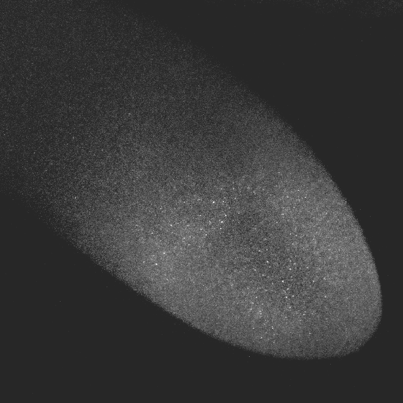

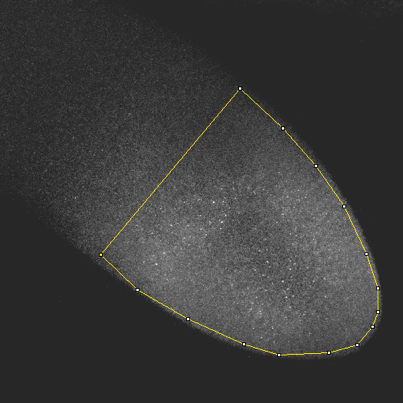

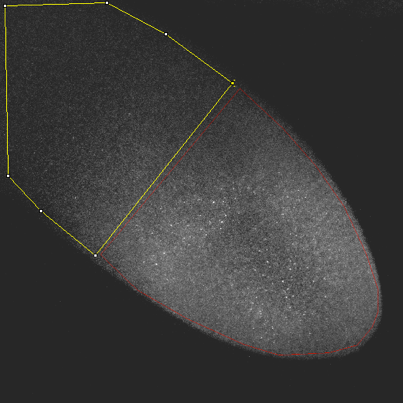

A	B	C

8. Repeat Steps 5 and 6 to obtain the average intensity of background noise.
9. Calculate the fold reduction of *lacZ* using the equation below.

$$Fold reduction=\frac{\left( hb-noise \right)_{I}}{\left( hb-noise \right)_{A}}$$

*hb* = average intensity of hunchback domain

*noise* = average intensity of background noise

I = measurements from embryos containing an inactive ribozyme

A = measurements from embryos containing an active ribozyme

F
