## Supplementary Table 2 for "Tunable self-cleaving ribozymes for modulating gene expression in eukaryotic systems"

| **Transfection conditions** | | |
| --- | --- | --- |
|  | 35mm plate | 24-well plate |
| Working volume (µl) | 2,000 | 500 |
| Cells seeded | 400,000 | 80,000 |
| DNA (ng) | 2,800 | 550 |
| DNA volume (µl) | 129 | 26 |
| FugeneHD reagent (µl) | 8 | 1.7 |
| Total volume added per plate/well (µl) | 125 | 25 |
